# Spatial Transcriptomics Inferred from Pathology Whole-Slide Images Links Tumor Heterogeneity to Survival in Breast and Lung Cancer

**DOI:** 10.1101/2020.07.02.183814

**Authors:** Alona Levy-Jurgenson, Xavier Tekpli, Vessela N. Kristensen, Zohar Yakhini

## Abstract

Digital analysis of pathology whole-slide images is fast becoming a game changer in cancer diagnosis and treatment. Specifically, deep learning methods have shown great potential to support pathology analysis, with recent studies identifying molecular traits that were not previously recognized on pathology H&E whole-slide images. Simultaneous to these developments, it is becoming increasingly evident that tumor heterogeneity is an important determinant of cancer prognosis and susceptibility to treatment, and should therefore play a role in the evolving practices of matching treatment protocols to patients. State of the art diagnostic procedures, however, do not provide automated methods for characterizing and/or quantifying tumor heterogeneity, certainly not in a spatial context. Further, existing methods for analyzing pathology whole-slide images from bulk measurements require many training samples and complex pipelines. Our work addresses these two challenges. First, we train deep learning models to spatially resolve bulk mRNA and miRNA expression levels on pathology whole-slide images (WSIs). Our models reach up to 0.95 AUC on held-out test sets from two cancer cohorts using a simple training pipeline and a small number of training samples. Using the inferred gene expression levels, we further develop a method to spatially characterize tumor heterogeneity. Specifically, we produce tumor molecular cartographies and heterogeneity maps of WSIs and formulate a heterogeneity index (HTI) that quantifies the level of heterogeneity within these maps. Applying our methods to breast and lung cancer slides, we show a significant statistical link between heterogeneity and survival. Our methods potentially open a new and accessible approach to investigating tumor heterogeneity and other spatial molecular properties and their link to clinical characteristics, including treatment susceptibility and survival.

## 1 Introduction

Digital pathology – the automated computer-vision analysis of pathology whole-slide images (WSIs) – is fast becoming a game changer in cancer diagnosis and treatment. Deep learning methods have been studied extensively in this context, and were recently shown to be efficient for certain tasks, such as detecting metastases [1, 2, 3], immune cells [4, 5, 6, 7], mitosis [8, 9] and tissue type [10, 11, 4] as well as for offering clinicians additional insights [12, 13, 14, 15]. In a recent study, [16] used deep learning to study immune geospatial variability and how it may affect the emergence of aggressive clinical phenotypes. These achievements led researchers to more recently explore whether such methods could go a step further, and identify molecular traits that are not known to be associated with cell/tissue morphology, such as mutations [17, 18], copy-number alterations [18, 19], gene expression [20, 18, 19] and hormone receptor status [15, 19, 21].

Simultaneous to these technological advances, the importance of tumor heterogeneity is being increasingly recognizes as a major feature associated with resistance to treatments and as a determinant of prognosis [22, 21, 23, 24, 25]. Polyclonality and tumor subclones – sub-populations of tumor cells that differ in molecular characteristics such as mutations, copy number aberrations and gene expression profiles – are a hallmark of cancer and may affect treatment outcome and disease progression [26, 27, 28, 7, 29, 30]. Bulk measurements, which offer high cell-coverage, have the potential to characterize tumor heterogeneity. A key downside, however, is that bulk measurements lack spatial context. Recent work showed that clonal estimation from single-region sampling is less accurate than that obtained by multi-region sampling and that single-region clonal composition estimations vary greatly between methods [31]. Realizing the importance of spatial context, new technologies for spatial transcriptomics have begun to emerge and are being increasingly used by the scientific community [32, 33, 34, 35, 36] alongside other methods to spatially resolve molecular measurements. In [32], spatial transcriptomics data, collected from 23 breast cancer patients, was used to train a deep neural network to predict spatial variation in gene expression. In a cohort of 41 gastric cancer patients, [21] discovered an association between heterogeneity and survival using a genome-wide single-nucleotide variation array to estimate the number of clones. This was followed by fluorescence in situ hybridization (FISH) to derive clonal locations for 3 samples across 4 target regions. [37] combined single cell RNA sequencing (scRNA-seq) with spatial transcriptomics to map and characterize the different cell populations in heterogeneous pancreatic tumors. In multiple myeloma, [38] inferred spatial organization from scRNA-seq data, using a clustering-based approach, to characterize immunological alterations occurring in the tumor microenvironment during disease progression. Others have used a computational approach to infer spatial position probabilities for individual cells from scRNA-seq data, enabling the spatial reconstruction of single-cell gene expression [39]. In [24], single-cell pathology subgroups were spatially resolved using mass cytometry imaging, covering an average of 2, 246 cells per image across 381 images, to characterize clonal populations in breast cancer. One of their findings associated a specific single-cell pathology subtype that comprised multiple epithelial cell communities with poor survival. In a similar setting, spatially-derived statistics from single-cell data were shown to improve prognostic predictions [23]. These findings emphasize the importance of characterizing tumor heterogeneity in a spatial context. However, single-cell and spatial transcriptomics techniques are expensive and are still limited in cell coverage compared to WSIs. While WSIs hold the potential to spatially resolve bulk molecular measurements using complete cell coverage [16, 20, 40, 18, 19], existing pipelines often require multiple modelling steps, large training sets and expert intervention. Furthermore, there is currently no method to automatically derive heterogeneity, both visually and quantitatively, from H&E WSIs.

In this paper, we present an automated method to both visualize and quantify tumor heterogeneity in the spatial context of WSIs using a simple pipeline, a small number of bulk-labeled training slides and no expert intervention. Applying our approach, we discover a significant link between tumor heterogeneity and survival outcome in breast and lung cancer. Briefly, we train deep neural networks to provide molecular cartographies of mRNA and miRNA expression from WSIs. We use a simple training-inclusion criteria to potentially reduce noise and facilitate model convergence speed and performance. We then use the inferred cartographies to produce heterogeneity maps and to quantify the level of heterogeneity within each WSI using our heterogeneity index (HTI). Applying our methods to breast and lung cancer slides, we show a significant link between heterogeneity and survival. An overview of our method is shown in Figure 1.

**Figure 1:**
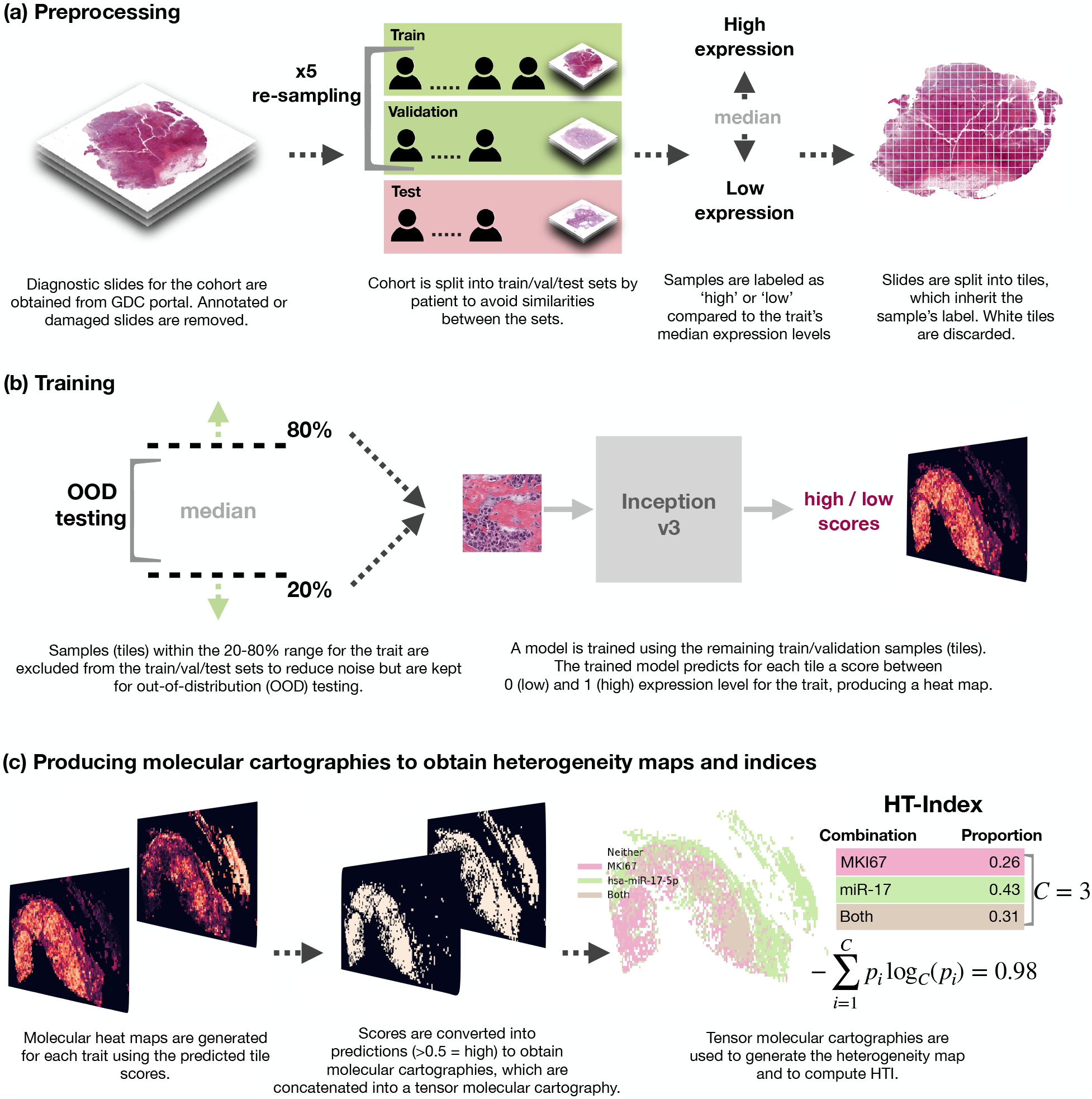
Overview of our methods: (a) prepossessing, (b) model training per trait and (c) producing tensor molecular cartographies per slide from which heterogeneity maps and indices are derived.

Our main contributions are: (1) A simple pipeline that maps from WSIs to gene expression levels, reaching up to 0.95 AUC on held-out random test sets. Our pipeline uses a single model architecture and requires only a small number of bulk-labeled training slides (with no expert annotations); (2) A method for constructing heterogeneity maps from the inferred gene-expression maps; (3) A heterogeneity index (HTI) that quantifies the level of heterogeneity within the WSI based on the spatial co-location of molecular traits; (4) A statistically significant link between tumor heterogeneity and survival outcome in two cancer cohorts. Our code is available online (Methods 4.5).

## 2 Results

### 2.1 Training models to identify gene expression on pathology whole-slide images

An overview of our methods is shown in Figure 1. We begin by obtaining formalin-fixed, paraffin-embedded (FFPE) hematoxylin and eosin (H&E) slides for TCGA BRCA and LUAD samples and matching mRNA and miRNA expression data (Methods 4.1). We discard any damaged, heavily stained or annotated slides as they can cause the model to focus on artifacts, including clues from pathologists’ markings. This results in 761 slides for breast and 469 for lung. We obtain matching normalized mRNA and miRNA expression data for a total of 10 molecular traits for breast and 5 for lung, as listed in Table 1. Breast mRNAs were chosen from the PAM50 genes [41] and lung mRNAs and all miRNAs (breast and lung) were based on the literature (see Methods 4.3 for further details).

**Table 1:** Results for breast and lung cohorts for test and OOD sets. Slides are scored using the percent positively classified tiles. Each of the 5 trained models is evaluated on the held-out test set and the two OOD sets separately. Average AUC results and range (brackets) are shown on the left, followed by ensemble AUC results for test, OOD-near and OOD-all. In bold are AUCs of at least 0.6.

|  | Trait | Per-slide AUC |  |  |  |
| --- | --- | --- | --- | --- | --- |
|  |  | Test<br>5-run average (range) | Test<br>ensemble | OOD-near<br>ensemble | OOD-all<br>ensemble |
| TCGA-BRCA | miR-17-5p | **0.83 (0.79-0.88)** | **0.87** | **0.62** | 0.56 |
|  | MKI67 | **0.74 (0.53-0.87)** | **0.87** | **0.72** | **0.63** |
|  | FOXA1 | **0.7 (0.6-0.76)** | **0.74** | **0.61** | 0.56 |
|  | MYC | **0.62 (0.58-0.67)** | **0.63** | 0.59 | 0.58 |
|  | miR-29a-3p | **0.62 (0.5-0.69)** | **0.67** | **0.65** | 0.55 |
|  | ESR1 | 0.58 (0.55-0.63) | 0.56 | **0.75** | 0.57 |
|  | CD24 | 0.57 (0.47-0.67) | 0.56 | 0.57 | 0.47 |
|  | FOXC1 | 0.53 (0.46-0.57) | 0.52 | **0.68** | **0.61** |
|  | ERBB2 | 0.5 (0.46-0.57) | 0.53 | 0.52 | 0.5 |
|  | EGFR | 0.5 (0.35-0.67) | 0.42 | 0.57 | 0.57 |
| TCGA-LUAD | miR-17-5p | **0.85 (0.72-0.95)** | **0.95** | 0.43 | 0.54 |
|  | KRAS | **0.64 (0.33-0.9)** | **0.73** | 0.56 | 0.52 |
|  | CD274 (PD-L1) | **0.61 (0.57-0.63)** | **0.6** | **0.64** | 0.51 |
|  | miR-21-5p | 0.55 (0.47-0.62) | 0.54 | **0.75** | **0.63** |
|  | EGFR | 0.44 (0.22-0.78) | 0.39 | 0.48 | 0.46 |

Each cohort is processed, trained and evaluated separately (Figure 1 (a) and (b)). We begin by randomly assigning subjects into a held-out test set (10%). We then randomly split the remaining subjects into train (80%) and validation (10%) sets 5 times (bootstrap sampling) to obtain 5 different and randomly selected train-validation sets. Note that these are further reduced in size by a simple inclusion criteria described below. We split on subjects rather than slides so that slides from the same subject are assigned to the same set to avoid similarities between test and train/validation. Importantly, the split is performed before any molecular trait is processed to ensure that all traits see the same split. We then assign each slide into one of the sets according to its subject’s association and proceed to label the slides and prepare our training set.

Per trait, we label each slide based on its sample’s expression percentile for that trait. We then split each slide into non-overlapping tiles of 512 × 512 at × 20 zoom, resulting in hundreds to thousands of tiles per slide (depending on the slide’s size). Following previous methods [17], each tile inherits its labels from its parent slide. We label each slide (and tiles) based on its percentile expression level: high (1) for those above median low (0) for those below. This raises the concern that not all tiles are representative of its slide label (e.g. a tile representing a healthy portion of a tumor sample), leading to noisy labeling. To potentially reduce noise and facilitate faster model convergence, we set aside complete slides (all corresponding tiles) with expression levels between the 20th and 80th percentiles and use them for out-of-distribution evaluation (Figure 1 (b)). These samples do not participate in the model’s train/validation/test sets. This inclusion cirteria enables us to efficiently train on a small number of slides (e.g. for BRCA, removing these slides after taking 80% for training leaves us with 760 × 0.8 × 0.4 = 243 training slides). We proceed to use the labeled tiles to train our models.

We train each trait separately and repeat this process 5 times (once for each of the random train/validation splits of the non-test samples). We use the Inception v3 classifier [42] as a single end-to-end model that predicts tile scores. This is the only model architecture used to obtain predictions. Additionally, no expert intervention is included at any step.

Once trained, we obtain the trait’s validation performances for all 5 runs by measuring the slide-level AUC for each of the five models (we evaluate using slide-level AUC following, e.g., [17, 43]). To do so, we first transition from predictions on tiles to predictions on slides by computing the slide’s percent positively classified tiles (as in [17]). We then compute the slide-level AUC for each model using these scores as the slide-level prediction. The final model for each of the 5 rounds uses the weights that had the best slide-level AUC on that round’s validation set.

Besides providing a better evaluation of our models’ performance, a key advantage of training multiple models on different train/validation splits is the option to combine them into an ensemble model [44]. We develop such an ensemble model for each trait by combining its top three models, as measured on their respective validation set, and taking their majority vote. For example, if the last three models from the 5 train/validation repeats yielded the best performances (each on their respective validation set) and their predictions for a given tile are: 0, 0, 1 then the ensemble will predict 0 for that tile by majority vote.

For each trait, we evaluate both its 5 models and its ensemble model on the trait’s held-out test set (test slides with expression levels within the 0-20% and 80-100% percentiles for that trait) as well as on its OOD sets (slides within the 20-80% percentiles). Neither of these test sets participated in the trait’s model development (train/validation) in any of the 5 rounds.

### 2.2 Achieving high prediction performance for identifying gene expression on pathology whole-slide images

We obtain high test performance rates across multiple traits. The number of slides in the test set varies across traits, depending on cohort (TCGA-LUAD is roughly half the size of TCGA-BRCA) and on label (expression) availability, and is therefore described per trait in brackets below. In Table 1, we observe that the average test performance measured across all 5 train/validation rounds (first column) obtains high AUCs for over half the traits. Most prominent are miR-17-5p (N=33), MKI67 (N=36) and FOXA1 (N=31) in breast as well as miR-17-5p (N=13) and KRAS (N=17) in lung. Especially noteworthy is miR-17-5p as it performed exceptionally well in both cohorts (for which model development and testing is completely separate), further suggesting that mRNA and miRNA expression can be detected from tissue morphology and that this may be applicable for other cancer types.

The use of ensemble models further improves the performance in nearly all traits. miR-17-5p improves from 0.83 to 0.87 AUC in breast and from 0.85 to 0.95 AUC in lung. We observe similar effects for MKI67 and FOXA1 in breast as well as for KRAS in lung. In Figure 2 a (breast) and d (lung) we can see the test-set distribution of the ensembles’ slide scores for the ground-truth labels across the different traits. We clearly observe that the images with bulk label above median have higher slide-level predictions. This is especially true for miR-17-5p (in both cohorts), MKI67, FOXA1, MYC, miR-29a-3p and ESR1 in breast as well as KRAS and CD274 (PDL-1) in lung. We also computed the correlation of our predicted levels to the actual percentiles (Supplementary S1). For example, in breast miR-17 we observe Spearman correlations of 0.63 (FDR p-value 8*e*^−04^), 0.21 (FDR p-value 0.02) and 0.13 (FDR p-value 0.01) in test, OOD-near and OOD-all respectively.

**Figure 2:**
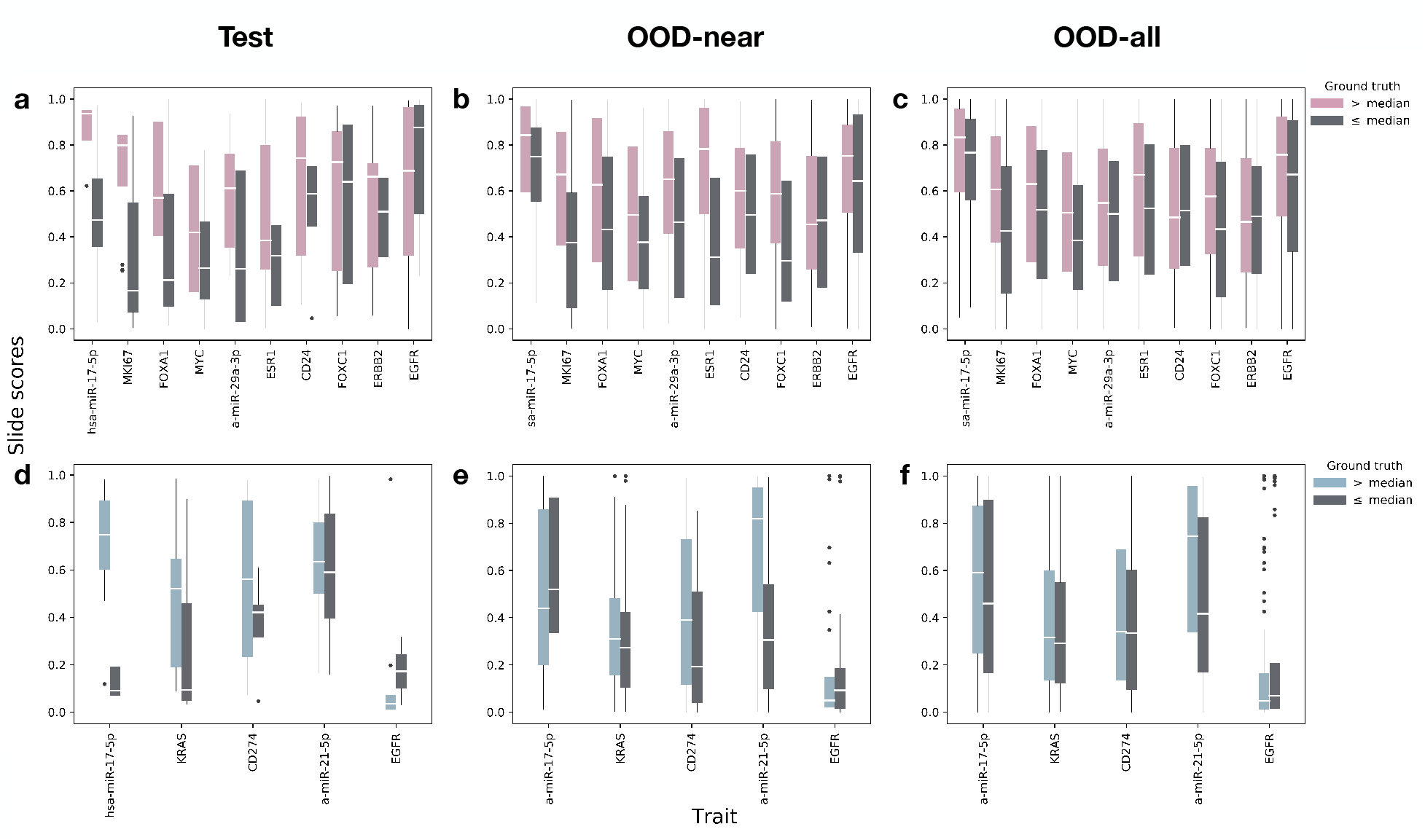
WSI score distribution for test, OOD-near and OOD-all sets compared to the actual bulk-measured values (lower values in gray). Scores are computed using the ensemble model for each trait (percent positive tiles by majority vote of the best 3). Middle white lines reflect the median slide score. Whiskers extend to 1.5 *× IQR*. (a)-(c) breast test, OOD-near and OOD-all respectively; (d)-(f) lung test, OOD-near and OOD-all (respectively). In the test set, especially worth noting are miR-17-5p in both cohorts, MKI67, FOXA1, MYC, mir-29a-3p and ESR1 in breast as well as KRAS and CD274 (PD-L1) in lung. In the OOD sets most notable are MKI67, FOXA1, MYC, ESR1 and FOXC1 in breast as well as miR-17-5p and miR-21-5p in lung.

To the best of our knowledge, these are the first models to automatically detect miRNA expression levels on H&E whole-slide images. Importantly, our method achieves these results using only a small number of training samples (243 slides for BRCA and 150 for lung, as explained in Section 2.1), a single model architecture and no expert knowledge. Model convergence took less than 12h (worst-case) on a single server with 8 low-cost GPUs (Tesla K80).

### 2.3 Performance for out-of-distribution (OOD) samples

An added advantage in setting aside the OOD slides is the ability to further challenge our models on out-of-distribution samples. We use the ensemble models for each trait. In Table 1 we report two types of AUC results on the OOD set: one for the next decile tier after the test set’s percentiles, i.e. 0.2-0.3 and 0.7-0.8, which we designate as near-distribution (OOD-near) and another for the full OOD set (OOD-all). As before, the number of slides available in the two OOD sets depends on the cohort and the intersection between slide and label availability for the trait. In the OOD-near set, breast has between 140 and 158 and lung has between 79 and 103 slides (roughly 20% from each cohort’s slides). The OOD-all set has between 423 and 454 slides for breast, and between 256 and 283 for lung (roughly 60% of each cohort’s slides - the remainder after using the top and bottom 20% for model development). We are able to test our models on such large test sets since, by design, none of the slides in their percentile range were included in the trait’s train/validation/test sets and therefore they were all held out. Our results demonstrate that the models extend to out-of-distribution samples relatively well. This can also be observed in Figure 2 b,c (breast) and e,f (lung) which depict the distribution of scores per label for the OOD-near and OOD-all sets. This is especially evident in the breast cancer cohort. For example, the score distributions for MKI67 for the “above median” (pink) class tend to be higher than those for the “below median” (gray) class. The overall better breast results are likely due to the larger amount of data available for training, emphasizing the importance of collecting larger data sets.

### 2.4 Tensor molecular cartographies of gene expression as a window to tumor heterogeneity

After confirming the performance of our models, we proceed towards our goal of analyzing tumor heterogeneiy (Figure 1 c). We do so by producing, for each slide, multiple molecular cartographies, each representing a single molecular trait. We then combine the molecular cartographies into a tensor molecular cartography to obtain a richer spatial representation of the molecular traits detected within the slide. Creating a molecular cartography for a single trait is straight forward: we simply spatially arrange the binary predictions of a slide’s tiles in a matrix so that the position of a tile’s score in the matrix corresponds to its position in the pre-tiled slide, as illustrated in (Figure 1 c left). Once we obtain several molecular cartographies for a single slide, we stack them into a tensor molecular cartography (Figure 1 c middle) that now represents the slide across several traits.

We use the tensor molecular cartography to visualize heterogeneity – this will later serve us to confirm our method for quantifying heterogeneity. As shown in Figure 1 (c right), we do so by identifying which tiles are positive for each possible combination of traits and assign different colors to each combination. For example, using two traits, A and B, we obtain a tensor of depth 2 (one molecular cartography per trait) and each tile is colored as one of three options: A (only), B (only) and Both. Figure 3 shows examples for such heterogeneity maps, along with their level of heterogeneity, as obtained using HTI described below. Such molecular maps of pathological images could be used by pathologists, in addition to routinely stained diagnostic markers, to further classify and identify cancer subtypes, potentially leading to better informed clinical decisions.

**Figure 3:**
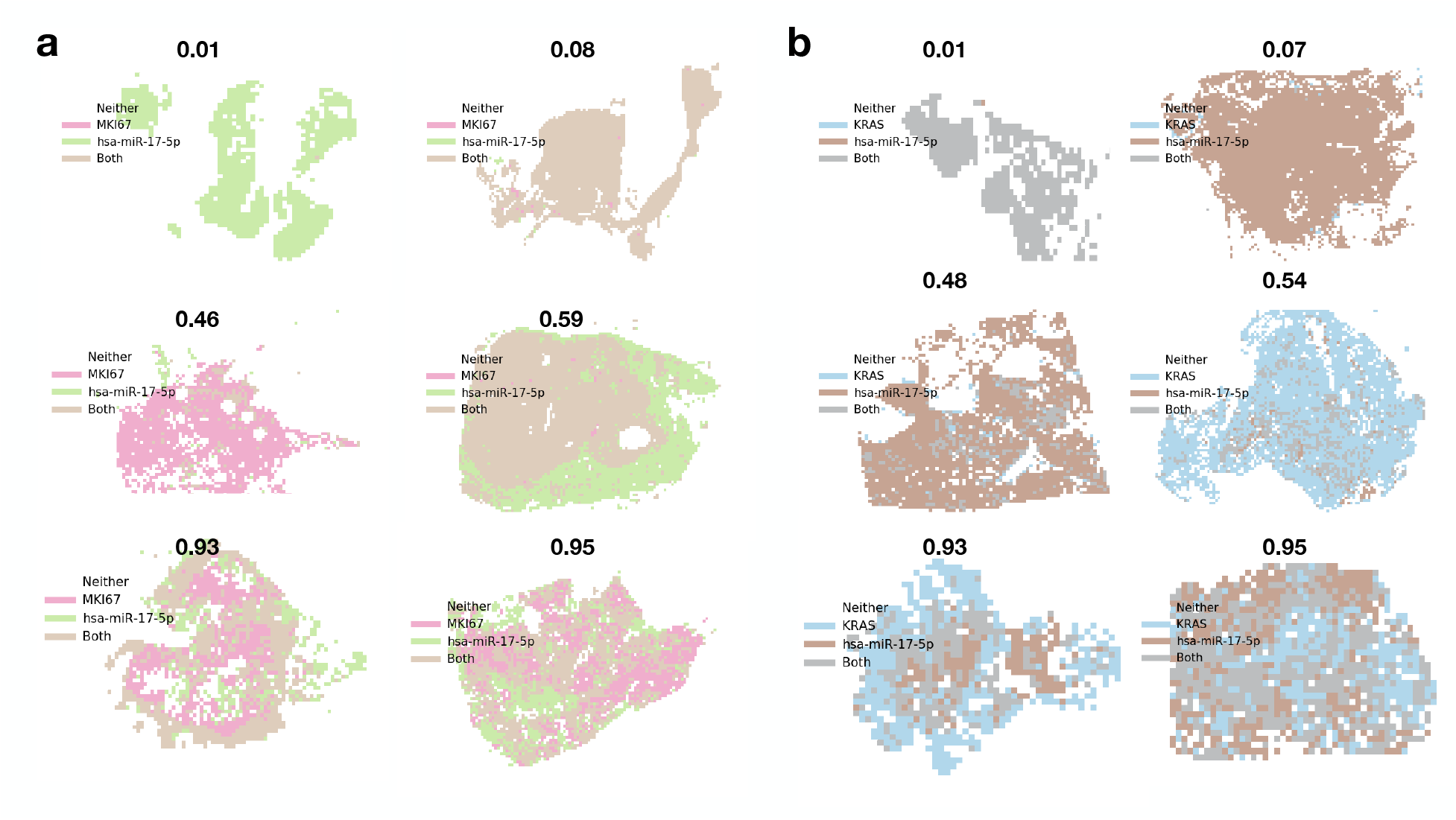
Heterogeneity maps and corresponding HTIs. (a) Breast cancer cohort with traits MKI67 (pink) and miR-17 (green). Brown indicates that both traits manifest (both models predicted positive). (b) Lung cancer cohort with traits KRAS (blue) and miR-17 (brown). Gray indicates both traits manifest. Corresponding HTIs appear directly above. Rows appear in increasing HTI order from top to bottom (top 0-0.1, middle 0.4-0.6, bottom 0.9-1).

### 2.5 Quantifying tumor heterogeneity from tensor molecular cartographies

As intra-tumor heterogeneity has been proposed as an obstacle to effective treatment and cancer eradication, we propose an approach to compute and quantify the level of heterogeneity in a given image from its tensor molecular cartography. We used a variation on Shannon’s entropy, commonly used to measure diversity and heterogeneity in various settings [45, 24]. Formally, we compute:

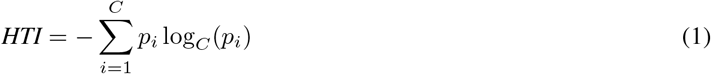

where *C* is the maximum number of non-empty trait combinations that may be observed on a slide and equals 2^| *traits*|^ − 1 (the number of subsets excluding the empty set), and *p*_*i*_ is the proportion of tiles for which exactly all models in combination *i* provided a positive prediction.

For example, given a tensor molecular cartography for two traits A and B (e.g. FOXA1 and MKI67), *C* = 3 (3 possible non-empty trait combinations: A (only), B (only) and Both). If the slide is homogeneous with nearly all of its tiles falling into one of these 3 options (say Both), then *p*_*Both*_ ≈ 1, *p*_*A*_ ≈ 0, *p*_*B*_ ≈ 0 (each of the single-trait molecular cartographies is nearly all 1s), resulting in an HTI of 0. If, however the slide is heterogeneous with 1*/*3rd of the tiles falling into each option then: *p*_*Both*_ = 1*/*3, *p*_*A*_ = 1*/*3, *p*_*B*_ = 1*/*3 and we obtain an HTI of 1. The logarithm base *C* guarantees that HTI ∈ [0, 1]. If A and B are two molecular traits (e.g. FOXA1 and MKI67), a high HTI would reflect there may be two subclones whereas a low HTI would reflect single clonal dominance.

We note that HTI can be applied to any set of binary matrices (or vectors) of identical shape, each of which describes the presence of a single trait. As such, it can be used in other settings involving the localization of clinically relevant traits. Figure 3 depicts several heterogeneity maps and their associated HTIs.

### 2.6 A statistically significant link between tumor heterogeneity and survival

We sought to understand whether tumor heterogeneity is linked to survival outcome and whether the link can be inferred through the spatial analysis of pathology whole-slide images. To do so, we combine the test and OOD slides and split them into high and low heterogeneity groups based on their HTIs. Specifically, for a given cohort, we first generate tensor molecular cartographies per slide using the ensemble models of the top two performing molecular traits for that cohort (from Table 1). We then compute HTI for each slide as described in Section 2.5 and split them into two groups: > 0.5 and ≤ 0.5 (high and low heterogeneity respectively). We perform survival analysis on these groups using Mantel’s log-rank test and Kaplan-Meier curves. We use only slides in the combined set of OOD and test slides. Since OOD slides are trait-dependent, we use only slides in the intersection of the two OOD sets.

Figure 4 describes the survival analysis results for each cohort using the top two traits by test performance from Table 1 in each (MKI67 and miR-17 for breast and miR-17 and KRAS for lung). In breast, high heterogeneity (blue) is significantly different from low heterogeneity (orange) with a log-rank p-value of 0.04 (Figure 4 (a)). In lung, we observe significant differences between higher and lower HTIs when heterogeneity differences are more distinct (> 0.7 vs ≤ 0.3), with a log-rank p-value of 0.07 (Figure 4 (c)).

**Figure 4:**
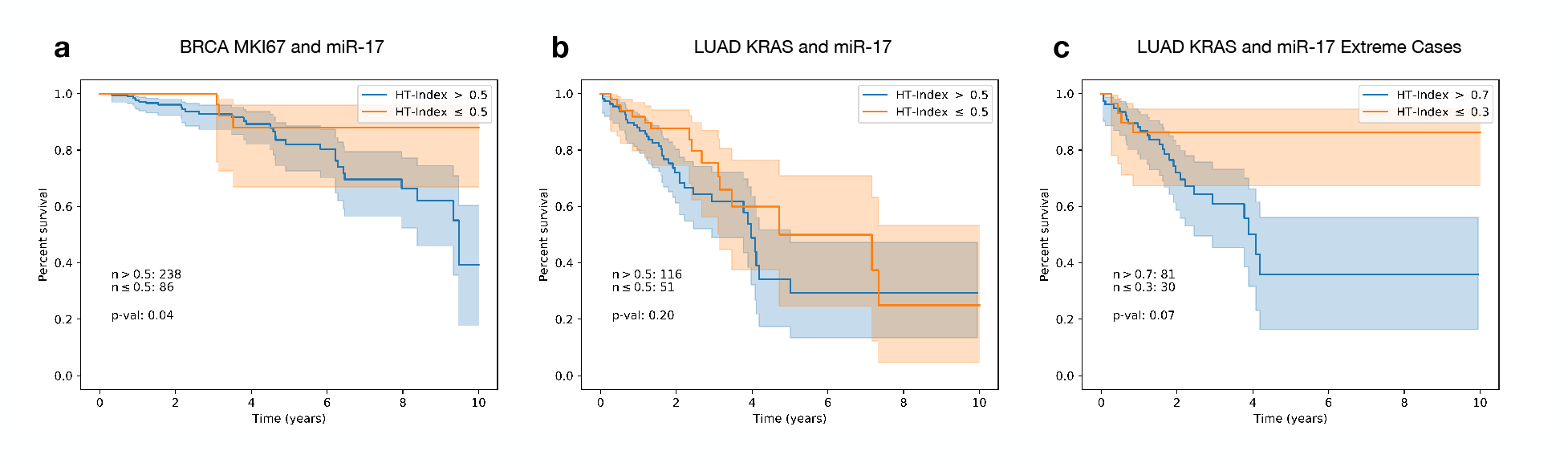
Survival analysis with respect to HTI derived from two traits in breast and lung cancer. For each trait combination in (a) and (b), slides were split into high and low HTI (> 0.5 and ≤ 0.5 or blue and orange respectively). In (c) slides were split into > 0.7 and ≤ 0.3 HTI. Each survival curve is shown with a 95% confidence interval.

## 3 Discussion

This work offers a method for analyzing tumor heterogeneity from the rich spatial data available in H&E WSIs. Using deep learning we created high resolution maps of multiple mRNA and miRNA expression levels within a whole-slide image and combined these maps into a tensor molecular cartography. We then used the tensor cartographies to spatially visualize and quantify tumor heterogeneity in the form of heterogeneity maps and HTI scores. While other methods, such as single-cell profiling and spatial transcriptomics, can also be used to infer heterogeneity, they are expensive, may lack sufficient spatial context and only cover a few thousand cells.

We applied our method to both breast and lung cancer pathology whole-slide images (H&E). We trained several models per trait and tested each of these models on both a held-out test set and two large out-of-distribution sets, containing hundreds of WSIs the models have never before encountered. Test results show that several mRNA and miRNA can be identified and localized automatically within whole-slide images with high AUCs. Furthermore, our results demonstrate this can be achieved using only a small number of training slides, a single model architecture and no expert intervention. Especially notable are our results for miR-17, which obtained high AUCs (up to 0.95) in both lung and breast. This is interesting in light of indications that miR-17 is over-expressed across many cancer types [46].

Using our models, we generated a tensor molecular cartography for each slide, enabling us to both visualize the distribution of traits, in the form of heterogeneity maps, and to compute HTI. By representing each patient with their HTI and performing survival analysis, we showed that high tumor heterogeneity is significantly linked to poor survival, especially in breast cancer. We stress that this link cannot be identified through obvious means by directly using expression levels (Figure 5) or through the PAM50 [41] breast cancer types, which are not associated with HTI (Figure 6). This analysis highlights the potential clinical value in producing tensor molecular cartographies and heterogeneity maps from H&E WSIs.

**Figure 5:**
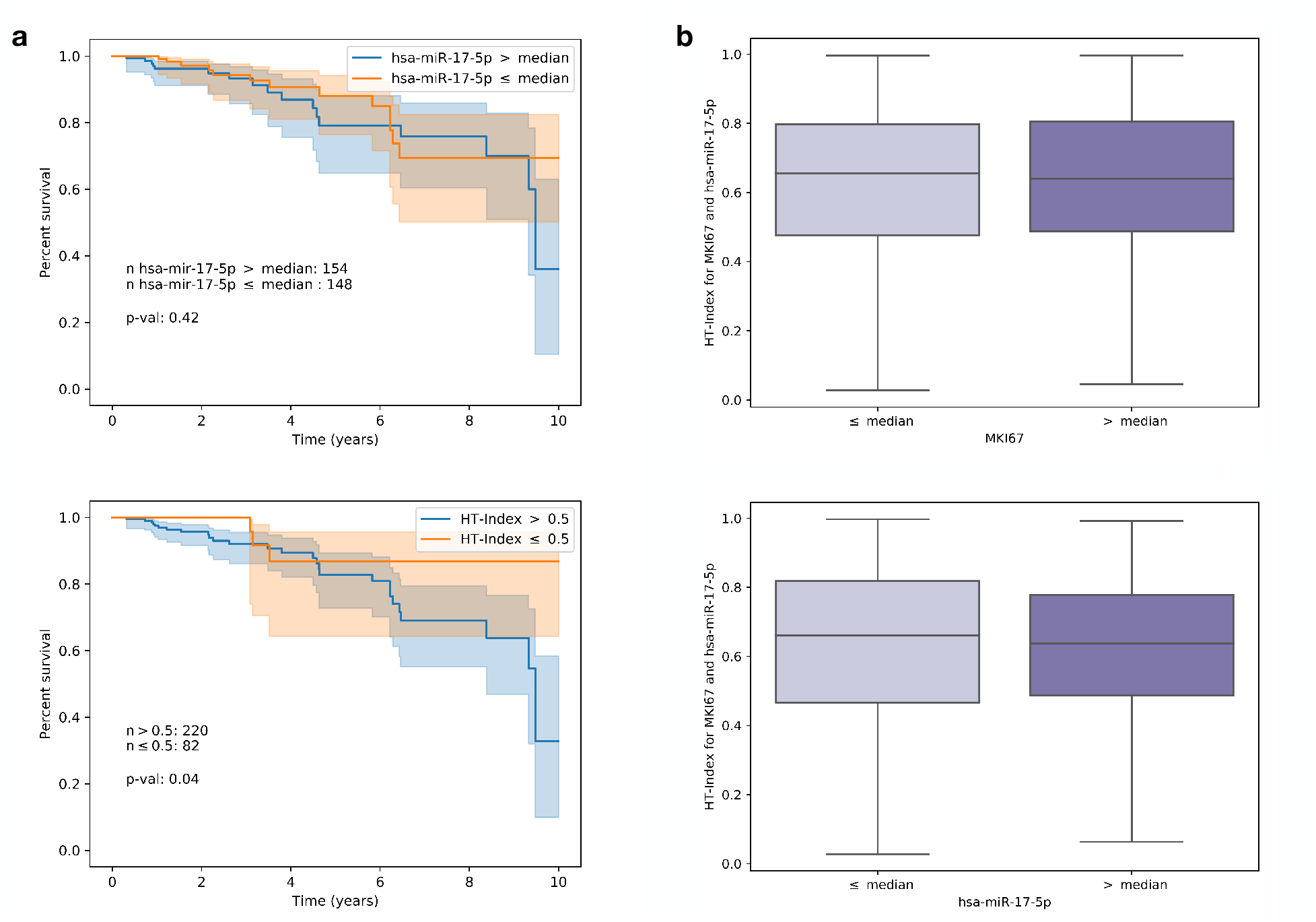
Left panel (a): Survival analysis using bulk-measured expression levels for miR-17 (top) compared to our approach (bottom) when applied to the same patient basis available for both. Right panel (b): distribution of HTI for above and below median ground-truth expression levels for MKI67 (top) and miR-17 (bottom).

**Figure 6:**
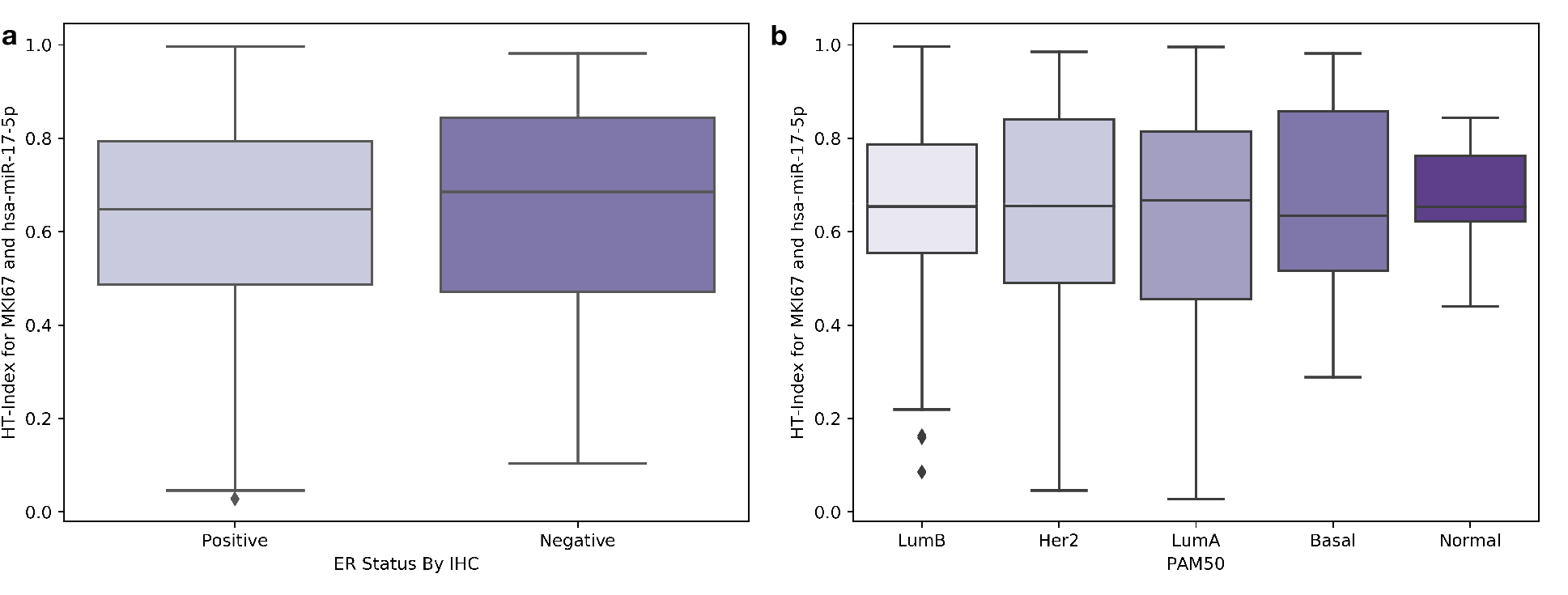
Distribution of HTI per ER status (a) and PAM50 type (b) in the data used for survival analysis.

Our methods open a window to further analyses. For example, from a molecular biology perspective, generating heterogeneity maps of miRNAs from the same family may be interesting in light of recent findings showing they are context (e.g. tissue) dependent [47]. Similarly, miRNA/mRNA relationships (as explored in [48, 49]) may be analyzed from a spatial perspective. From a technical aspect, it may be interesting to explore whether transfer learning between molecular traits or cohorts is possible and to what extent. Also a multilabel approach may be possible, although it may require larger datasets and careful label-balancing to obtain satisfactory results across all traits.

As heterogeneity plays an increasingly key role in cancer treatment, providing researchers and practitioners with a solution to view the distribution of clinically relevant traits that are not currently visible on slides may be of great value. Since H&E slides are a standard component of routine diagnostic protocols, a natural solution may be offered by the fast and automated digital mapping of the molecular landscape within H&E slides. One such solution is offered through our simple pipeline for producing tensor molecular cartographies and heterogeneity maps from WSIs.

As our approach requires a single model architecture, uses a small number of training slides and does not call for expert annotations, it can apply to many more traits and in broader contexts. As a future direction we hope to combine our approach with single-cell data and spatially resolved transcriptomics data to obtain finer resolution mapping, improving the relevance of both H&E and transcriptomics. Molecular cartography from H&E potentially enables a new approach to investigating tumor heterogeneity and other spatial molecular properties and their link to clinical characteristics. An interesting aspect of such endeavors will be to link spatial properties to treatment susceptibility and precision care.

## 4 Methods

### 4.1 Data

All whole-slide images are available online at the GDC repository (https://portal.gdc.cancer.gov/repository), by selecting *Diagnostic Slide* under *Experimental Strategy* for the relevant project (e.g. TCGA-BRCA). Matching expression levels were obtained from the GDC’s website at: https://gdc.cancer.gov/about-data/publications/pancanatlas (see RNA and miRNA). Matching survival data were obtained from cBioportal at: http://www.cbioportal.org/study/clinicalData from the following files: “Breast Invasive Carcinoma (TCGA, Firehose Legacy)” and “Lung Adenocarcinoma (TCGA, Firehose Legacy)”.

### 4.2 Training

All of our models were developed in TensorFlow [50] using the Inception v3 classifier [42] with the last layer modified to one output. All models are trained from random initialization, following previous work showing improved performance by fully training Inception v3 [17]. Each of the 5 train/validation rounds is trained on mini-batches of labeled tiles from the training set using the Adam optimizer [51]. Models were evaluated on the labeled validation tiles every 1/16th epoch (full pass on all training tiles) to avoid overfitting caused by tile similarities between mini-batches (each slide contains hundreds to thousands of tiles, many of which are likely to be similar to one another). No data augmentations were performed, except for random horizontal and vertical flips of the training tiles to further reduce overfitting. Learning rate started at 0.001 and was decayed when performance on the validation set plateaued for 10 steps, with early stopping after 30 steps of no validation improvement. Each final trained model used the weights that performed best on its validation set.

To potentially reduce noise and facilitate faster model convergence, we set aside complete slides (all corresponding tiles) with expression levels between the 20th and 80th percentiles. These were used for out-of-distribution evaluation and did not participate in the model’s train/validation/test sets. This enabled us to efficiently train on a small number of slides. For example, for BRCA, after removing these slides, and taking 80% for training, we were left with 760 × 0.4 × 0.8 = 243 training slides.

Each model was trained on a single Tesla K80 machine with 8 GPUs and took at most 12 hours until convergence on the validation set (when the aforementioned early stopping was invoked). Mini-batch size per GPU replica was 18, for a total of 18 × 8 = 144 tiles per training step.

### 4.3 mRNA and miRNA selection

#### mRNAs

For breast, we sorted PAM50 [41] genes by their expression variance in our dataset and selected from the top (highest variance). CD24 is not in PAM50, but is one of the highest expression-varying mRNAs in the cohort and is over-expressed in many cancers [52]. For lung, we based our selection on previous research, e.g. [53, 17, 54, 55].

#### miRNAs

Selection of miRs is based on previous work identifying miR-17 and miR-29 among the top up-regulated and down-regulated miRNAs (respectively) in breast cancer [49]. Others have also associated miR-17 [56, 57, 58] and miR-29 [59, 60] with breast cancer. miR-17 was also chosen for lung since the miR-17 family was shown to be universally over-expressed in many cancers, including lung [46], and has been directly associated with lung cancer [61, 62, 63] as has miR-21 [64, 65].

### 4.4 Survival analysis

Each patient is represented by a single HTI to perform survival analysis. Where patients were associated with more than one whole-slide image (e.g. a patient with diagnostic slides DX1 and DX2), the DX1 slide was used to determine HTI. For data used for survival see Methods 4.1. Analysis was performed using the Lifelines package for Python (https://lifelines.readthedocs.io/en/latest/).

### 4.5 Code availability

The code used for this work is publicly available under: https://github.com/alonalj/PathoMCH.

## 4.6 Author contribution

ALJ and ZY designed the study and developed the methods. ALJ performed all data analysis. XT and VK performed follow up analysis. XT curated data and prepared it for analysis. ZY supervised the study. ALJ and ZY wrote the manuscript with contributions from all authors.

## 4.7 Acknowledgement

Cloud computation for this project was partially funded by Google Cloud Platform. We thank the Technion Computer Science Faculty for generously supporting ALJ. This project received funding from the European Union’s Horizon 2020 Research and Innovation Programme under Grant agreement No. 847912. We thank the Yakhini Research Group for important discussions and input. We thank Doron Lipson for critical reading and important suggestions. We thank Øystein Garred for helpful discussions and for clarifying important pathology related aspects.

