## Supplementary for "Spatial Transcriptomics Inferred from Pathology Whole-Slide Images Links Tumor Heterogeneity to Survival in Breast and Lung Cancer"

### 1 Supplementary Material

#### 1.1 Supplementary Note S1

Table S1: Spearman correlations and FDR-corrected p-values for breast and lung cohorts across test, OOD-near and OOD-all data sets. Slides are scored using the percent positively classified tiles produced by the ensemble model for each trait. Ground-truth values are the actual slide percentiles (0.1, 0.2 etc.). In bold are significant FDR-corrected p-values $\leq 0.1$.

| | **Gene / Trait** | **Test** | | **OOD-near** | | **OOD-all** | |
|---|---|---|---|---|---|---|---|
| | | **Spearman** | **p-value (FDR)** | **Spearman** | **p-value (FDR)** | **Spearman** | **p-value (FDR)** |
| TCGA-BRCA | miR-17-5p | **0.63** | **8e-04** | **0.21** | **0.02** | **0.13** | **0.01** |
| | MKI67 | **0.58** | **1e-03** | **0.38** | **4e-06** | **0.27** | **6e-08** |
| | FOXA1 | **0.44** | **0.04** | **0.19** | **0.03** | **0.12** | **0.01** |
| | MYC | 0.36 | 0.11 | **0.16** | **0.07** | **0.13** | **0.01** |
| | miR-29a-3p | 0.38 | 0.11 | **0.27** | **4e-03** | **0.13** | **0.01** |
| | ESR1 | 0 | 1.00 | **0.43** | **5e-07** | **0.21** | **3e-05** |
| | CD24 | 0.10 | 0.93 | 0.12 | 0.17 | 8e-03 | 0.86 |
| | FOXC1 | 4e-03 | 1.00 | **0.31** | **3e-04** | **0.21** | **3e-05** |
| | ERBB2 | 0.07 | 0.99 | 0.03 | 0.70 | 0.02 | 0.72 |
| | EGFR | 0.05 | 1.00 | 0.12 | 0.16 | **0.12** | **0.02** |
| TCGA-LUAD | miR-17-5p | 0.56 | 0.22 | -1e-01 | 0.40 | 1e-03 | 0.98 |
| | KRAS | 0.38 | 0.32 | 0.11 | 0.40 | 0.03 | 0.74 |
| | CD274 (PD-L1) | 0.39 | 0.39 | **0.23** | **0.10** | 0.07 | 0.43 |
| | miR-21-5p | 0.17 | 0.49 | **0.43** | **2e-04** | **0.24** | **4e-04** |
| | EGFR | 0.27 | 0.49 | -3e-02 | 0.78 | -7e-02 | 0.43 |
